# A distributed residue network permits conformational binding specificity in a conserved family of actin remodelers

**DOI:** 10.1101/2021.05.27.445944

**Authors:** Theresa Hwang, Sara S Parker, Samantha M. Hill, Meucci W. Ilunga, Robert A. Grant, Ghassan Mouneimne, Amy E. Keating

## Abstract

Metazoan proteomes contain many paralogous proteins that have evolved distinct functions. The Ena/VASP family of actin regulators consists of three members that share an EVH1 interaction domain with a 100% conserved binding site. A proteome-wide screen revealed ciliary protein PCARE as a high-affinity ligand for ENAH EVH1. Here we report the surprising observation that PCARE is ~100-fold specific for ENAH over paralogs VASP and EVL and can selectively bind and inhibit ENAH-dependent adhesion in cells. Specificity arises from a mechanism whereby PCARE stabilizes a conformation of the ENAH EVH1 domain that is inaccessible to family members VASP and EVL. Structure-based modeling rapidly identified seven residues distributed throughout EVL that are sufficient to differentiate binding by ENAH vs. EVL. By exploiting the ENAH-specific conformation, we rationally designed the tightest and most selective ENAH binder to date. Our work uncovers a conformational mechanism of interaction specificity that distinguishes highly similar paralogs and establishes tools for dissecting specific Ena/VASP functions in processes including cancer cell invasion.

## Introduction

Metazoan signal transduction networks have evolved a high degree of complexity using adapter proteins that are specialized to make many interactions and/or highly specific interactions (Rowland et al., 2017; Zarrinpar et al., 2003). Signaling complexity arises in part from a plethora of interaction domain families such as the SH3, SH2, and PDZ domains. The facile recombination and insertion of modular domains to generate diverse protein architectures have enabled the evolution of new signaling circuits (Pawson and Nash, 2000).

Many modular interaction domains bind to short linear motifs (SLiMs), which occur as stretches of 3-10 consecutive amino acids in intrinsically disordered regions of proteins. The SLiM-binding specificity profiles of different paralogous members in a family of domains are often highly overlapping, yet individual members can in some cases engage in highly selective interactions (Xin et al., 2014; Hause et al., 2012). For example, a SLiM in Pbs2 binds only the SH3 domain of Sho1 out of the 27 SH3 domains in yeast to activate the high-osmolarity stress response pathway (Zarrinpar et al., 2003). Conversely, the actin assembly protein Las17 binds promiscuously to many SH3 domains, including Sho1, to drive actin-related processes such as endocytosis in yeast (Kelil et al., 2016; Robertson et al., 2009).

The Ena/VASP proteins are a family of actin regulators involved in functions ranging from T-cell activation to axon guidance (Kwiatkowski et al., 2003). There are three paralogs in mammals: ENAH, VASP, and EVL. All three proteins contain an N-terminal EVH1 interaction domain responsible for subcellular localization and a C-terminal EVH2 domain that polymerizes actin. The two domains are connected by a linker, predicted to be largely disordered, that contains binding motifs for other proteins. The EVH1 domain binds SLiMs with the consensus motif [FWYL]PXΦP, where X is any amino acid and Φ is any hydrophobic residue (Ball et al., 2000). This sequence, referred to here as the FP4 motif, binds the EVH1 domain as a polyproline type II (PPII) helix.

Ena/VASP proteins have evolved both overlapping and paralog-specific cellular functions. On the one hand, ENAH, VASP, and EVL can all bind to FP4 motifs in lamellipodin to promote actin assembly at the leading edge (Krause et al., 2004; Hansen and Mullins, 2015). Single deletions of Ena/VASP paralogs lead to mild phenotypic defects in mice, indicating that the paralogs can functionally compensate for each other, whereas triple mutant mice die after proceeding to late embryogenesis (Aszódi et al., 1999; Lanier et al., 1999; Kwiatowski et al., 2007). On the other hand, the three paralogs participate in distinct pathways. ENAH, alone, promotes haptotaxis of breast cancer cells through fibronectin gradients and regulates translation of specific mRNAs in developing axons, implicating it in functions beyond its role in actin polymerization (Oudin et al., 2016; Vidaki et al., 2018). In addition, whereas ENAH and VASP promote the invasive potential of migratory breast cancer cells, EVL suppresses breast cancer invasion (Roussos et al., 2011; Zhang et al., 2009; Mouneimne et al., 2018).

The FP4-binding pocket of the EVH1 domain is 100% conserved across ENAH, VASP, and EVL, and the paralogs share 62-72% sequence identity over the entire domain (Figure 1A, B). Consequently, the three EVH1 domains recognize many common binding partners. Nevertheless, some proteins bind selectively to certain paralogs. The LIM3 domain of testin binds specifically to the ENAH EVH1 domain in a region adjacent to the highly conserved FP4-binding pocket; this is the only example of an endogenous Ena/VASP EVH1 binding partner where the mechanistic basis for specificity is defined (Boëda et al., 2007). Testin binds specifically because it contacts residues that have diverged across paralogs to form distinct surfaces, which is a common mechanism for achieving specificity (Skerker et al., 2008; Bardwell et al., 2009; Schreiber and Keating, 2011).

**Figure 1.**
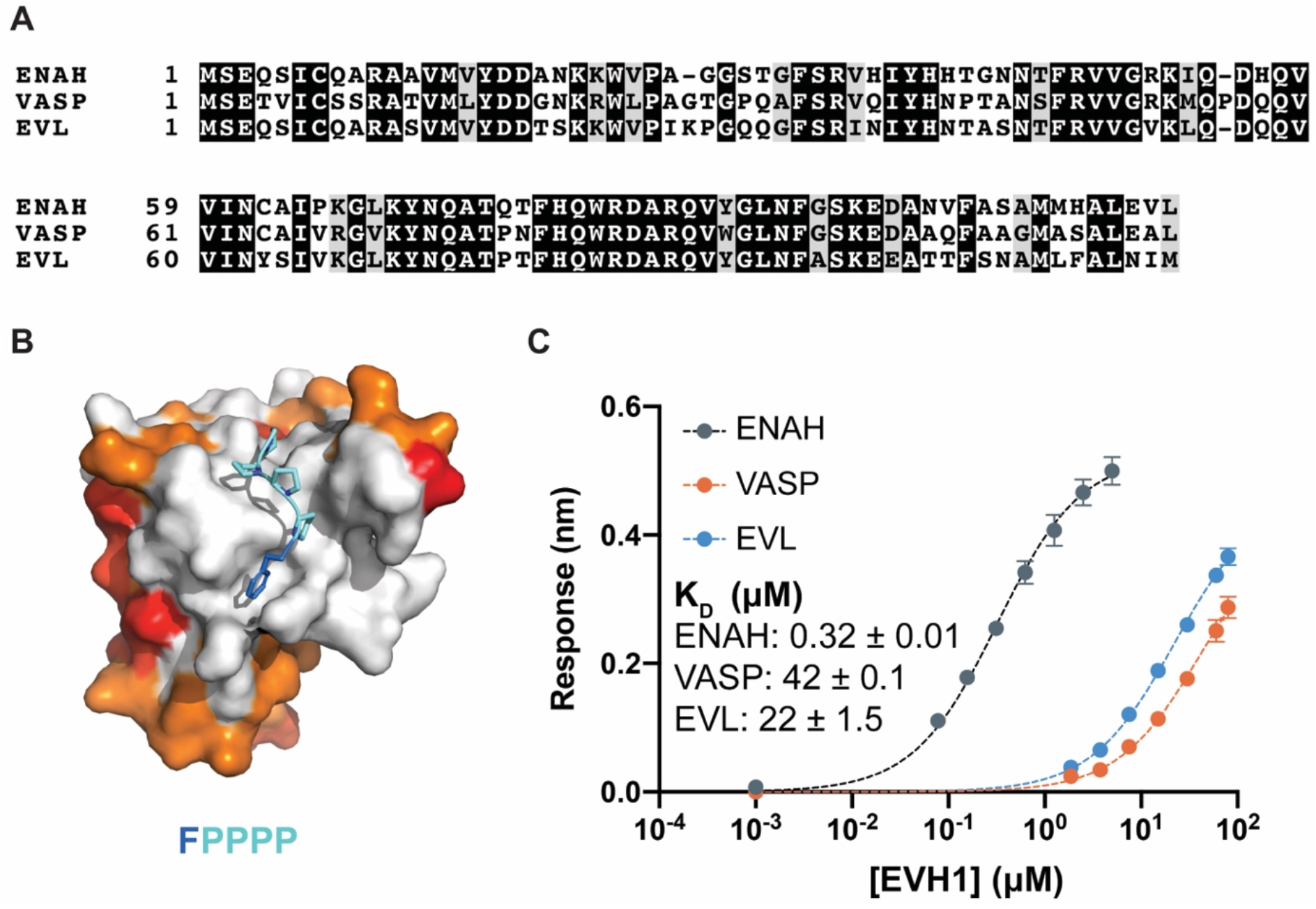
Ena/VASP family EVH1 domains are highly conserved but differ in their affinity for PCARE B. (A) Sequence alignment of the EVH1 domains of ENAH, VASP, and EVL. Black denotes residues that are 100% conserved; residues in gray share similar physiochemical properties. (B) Surface representation of ENAH EVH1 bound to an FP4 peptide (PDB 1EVH) highlighting conservation among Ena/VASP paralogs. Residues shared between all three paralogs are white; residues shared by two paralogs are orange, residues that differ in each paralog are red. (C) Biolayer interferometry data and curves fit to determine the dissociation constants for PCARE B binding to ENAH, EVL, or VASP EVH1 domains. Error reported as the standard deviation of two replicates.

Using a high-throughput screen of the human peptidome (Hwang et al. co-submission), we identified, among other interaction partners, a peptide from ciliary protein PCARE that binds with high affinity to ENAH EVH1 domain. Here we report that PCARE is selective for binding to ENAH in preference to EVL or VASP, despite containing an FP4 motif that can engage the perfectly conserved FP4-binding site. We demonstrate that this extended PCARE peptide can be used as a tool to perturb ENAH-specific interactions in cells. We also reveal the mechanistic basis behind the surprising paralog selectivity, which involves an epistatic residue network in the ENAH EVH1 domain that allows ENAH to adopt a conformation that is not accessible to paralogs VASP and EVL. An alpha-helical extension C-terminal to the FP4 motif in PCARE engages and stabilizes this ENAH-specific conformation using a noncanonical binding mode. These observations reveal a strategy to obtain binding specificity in a highly conserved family that must also make promiscuous interactions. Finally, we demonstrate how information about PCARE binding can be leveraged to design synthetic peptides with further-enhanced and unprecedented affinity and specificity for ENAH.

## Results

### An extended SLiM from ciliary protein PCARE binds selectively to ENAH EVH1 and reduces adhesion in mammalian cells

In a separate study, we performed a proteomic screen that identified peptides that bind to the ENAH EVH1 domain with dissociation constants (K_D_) primarily in the low- to mid-micromolar range (2 μM – 60 μM). The highest affinity hit from our screen was a 36-residue peptide from ciliary protein PCARE (PCARE^813-848^) that bound to ENAH with a K_D_ of 0.19 μM (Hwang et al., 2021). Truncation studies showed that 23-residue PCARE^826-848^, which we call PCARE B, maintains high affinity for ENAH EVH1 (K_D_ = 0.32 μM). To explore the paralog specificity of PCARE B, we quantified binding to EVL and VASP EVH1 domains using biolayer interferometry and discovered a 70-140-fold preference for binding to ENAH over the EVL or VASP (Figure 1C). This was initially very surprising, given that PCARE B includes the FP4 sequence LPPPPP and the Ena/VASP paralogs are 100% conserved in the FP4-binding groove (Figure 1B). However, this was also an exciting observation, because it suggested that PCARE B could serve as a reagent for selectively targeting ENAH in cell biological studies of its functions.

To test for association of Ena/VASP proteins and PCARE B in cells, we used Ena/VASP-family-deficient cell line MV^D7^ (Bear et al., 2000) and expressed the individual paralogs ENAH, EVL, and VASP to evaluate their interactions with PCARE. MV^D7^ are embryonic fibroblasts derived from *ENAH^-/-^ VASP^-/-^* mice, with low *EVL* expression (Damiano-Guercio et al., 2020). We used an shRNA against *EVL* to further decrease residual EVL (Figure S1). We tagged PCARE B with mRuby2 (mRuby2-PCARE B) and co-expressed this construct with GFP, or GFP fusions to ENAH, an EVH1 deletion mutant of ENAH (ΔEVH1-ENAH), EVL, or VASP. ENAH, EVL, and VASP all robustly localized to focal adhesions as previously observed (Puleo et al., 2019) while ΔEVH1-ENAH localization was cytoplasmic. mRuby2-PCARE B exhibited diffuse cytoplasmic localization under all conditions except when co-expressed with ENAH, in which case mRuby2-PCARE B was moderately enriched at focal adhesions, consistent with ENAH-PCARE B interaction (Figure 2A). Plots of fluorescence intensity along a line passing through focal adhesions show co-localization of PCARE B and ENAH (Figure 2B).

**Figure 2.**
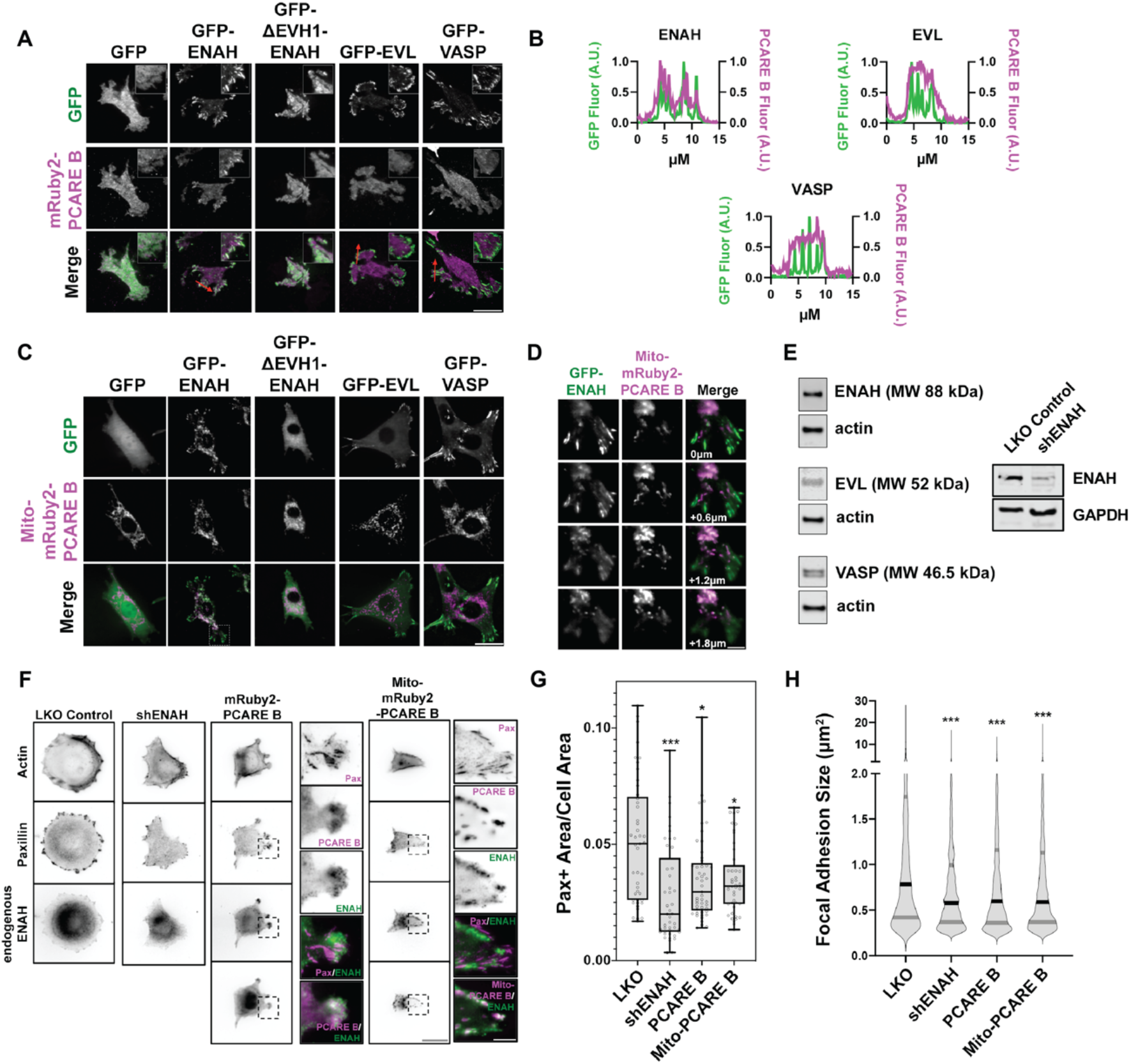
A PCARE-derived peptide selectively recruits ENAH in cells. (A) Live MV^D7^ shEVL cells expressing mRuby2-PCARE B with GFP, GFP-ENAH, GFP-dEVH1-ENAH, GFP-EVL, or GFP-VASP, imaged using TIRF microscopy. Scale bar = 25 μm. (B) Normalized fluorescence intensity of GFP signal (left axis) and mRuby2-PCARE B signal (right axis) along a line drawn through focal adhesions, indicated by the red arrow in (A). (C) Live MV^D7^_shEVL_ cells expressing Mito-mRuby2-PCARE B with GFP, GFP-ENAH, GFP-dEVH1-ENAH, GFP-EVL, or GFP-VASP. Image is a maximum intensity projection of z-stack acquired using widefield fluorescence microscopy. Scale bar = 25 μm. (D) Magnified region of interest indicated by the box in (C), showing co-localization of Mito-mRuby2-PCARE B and GFP-ENAH at increasing depths in the cell. Scale bar = 5 μm. (E) At left, representative western blot showing expression of ENAH, EVL, and VASP in MCF7 cells. At right, representative western blot showing the magnitude of knockdown using *ENAH*-targeted shRNA. (F) Immunofluorescence labeling of MCF7 cells expressing non-targeting LKO control, *ENAH*-targeting shRNA, mRuby2-PCARE B, or Mito-mRuby2-PCARE B. Cells were fixed eight hours after plating and immunolabeled for focal adhesion marker paxillin and endogenous ENAH, and additionally stained with phalloidin for Factin. Box indicates positions of magnified regions of interest (ROI). Scale bar = 25 μm, magnified ROI scale bar = 5 μm. (G) Box-and-whisker plot of total paxillin-positive area per cell normalized to the total cell area, for indicated conditions. N= 3 biological replicates, n= 45-49 cells. (H) Violin plot of individual focal adhesion size (for adhesions greater than 0.25 μm^2^) for indicated conditions. The central black line indicates the median, peripheral gray lines indicate interquartile ranges. N = 3 biological replicates, n = 1452-2159 individual adhesions.

Historically, a mito-tagged ActA peptide has served as a valuable sideways knockout tool that can deplete Ena/VASP proteins from other cytoplasmic locations by recruiting them to the mitochondria. This strategy has been used to dissect Ena/VASP-dependent functions (Bear et al., 2000). To test whether PCARE B could serve a similar function, but with specificity for ENAH, we tagged PCARE B with a mitochondrial localization sequence, generating Mito-mRuby2-PCARE. In MV^D7^_shEVL_ cells with exogenously expressed Ena/VASP paralogs, ENAH was strongly colocalized with Mito-mRuby2-PCARE B at mitochondria, in addition to being enriched at focal adhesions, while EVL and VASP remained entirely localized to focal adhesions and did not localize to mitochondria (Figure 2C, D). Together, these findings indicate that PCARE B sequesters ENAH, but not EVL or VASP, to artificial localization sites such as the mitochondria, supporting interaction specificity between PCARE B and ENAH in cells.

We also tested whether cytoplasmic expression of PCARE B could disrupt the function of endogenous ENAH. We examined focal adhesion maturation as a readout of ENAH function in MCF7 breast cancer cells, which express significant levels of ENAH (Figure 2E). We compared the effects of PCARE B expression to the effects of ENAH knockdown by shRNA by assessing cell adhesion, using paxillin immunofluorescence labeling to delineate focal adhesions (Figure 2F). As expected, ENAH knockdown was associated with diminished adhesion and smaller focal adhesion size as compared to cells expressing non-targeting shRNA (Figures 2F-H). Intriguingly, expression of PCARE B resulted in a similar decrease in adhesion, suggesting suppression of ENAH function at focal adhesions (Figures 2F-H). Importantly, the expression of PCARE B in MCF7 cells shifted the enrichment of ENAH from focal adhesions to membrane protrusions. ENAH enrichment at protrusions was not observed in MCF7 cells expressing Mito-PCARE B, which exhibited strong mitochondrial localization of ENAH and markedly decreased adhesion. This suggests that in MCF7 cells, blockade of the EVH1 domain by cytosolic PCARE B liberates ENAH from focal adhesions while permitting other EVH1-independent interactions elsewhere in the cell. In contrast, Mito-PCARE B serves to sink ENAH away from its normal sites of action at focal adhesions and the cell membrane.

### FP4 motif-flanking elements in ciliary protein PCARE confer high affinity by inducing noncanonical binding

To understand the structural basis for the high-affinity and selective interaction of PCARE B with ENAH, we solved a crystal structure of ENAH EVH1 domain fused to a 36-mer PCARE sequence to 1.65 Å resolution. Twenty-one residues of the 36-mer PCARE peptide were fully resolved in the electron density (PCARE^828-848^, Figure 3A) and led to the surprising discovery that the LPPPP motif in PCARE binds at the expected canonical site, but in the opposite orientation from previously observed Ena/VASP EVH1 domains engaged with proline-rich peptides (Ball et al., 2000; Prehoda et al., 1999; Federov et al., 1999) (Figure 3A, B). PCARE^828-848^ uses a 14-residue alpha helix-rich extension C-terminal to the LPPPP motif to make additional contacts to an extended region on the EVH1 domain, explaining its high affinity.

**Figure 3.**
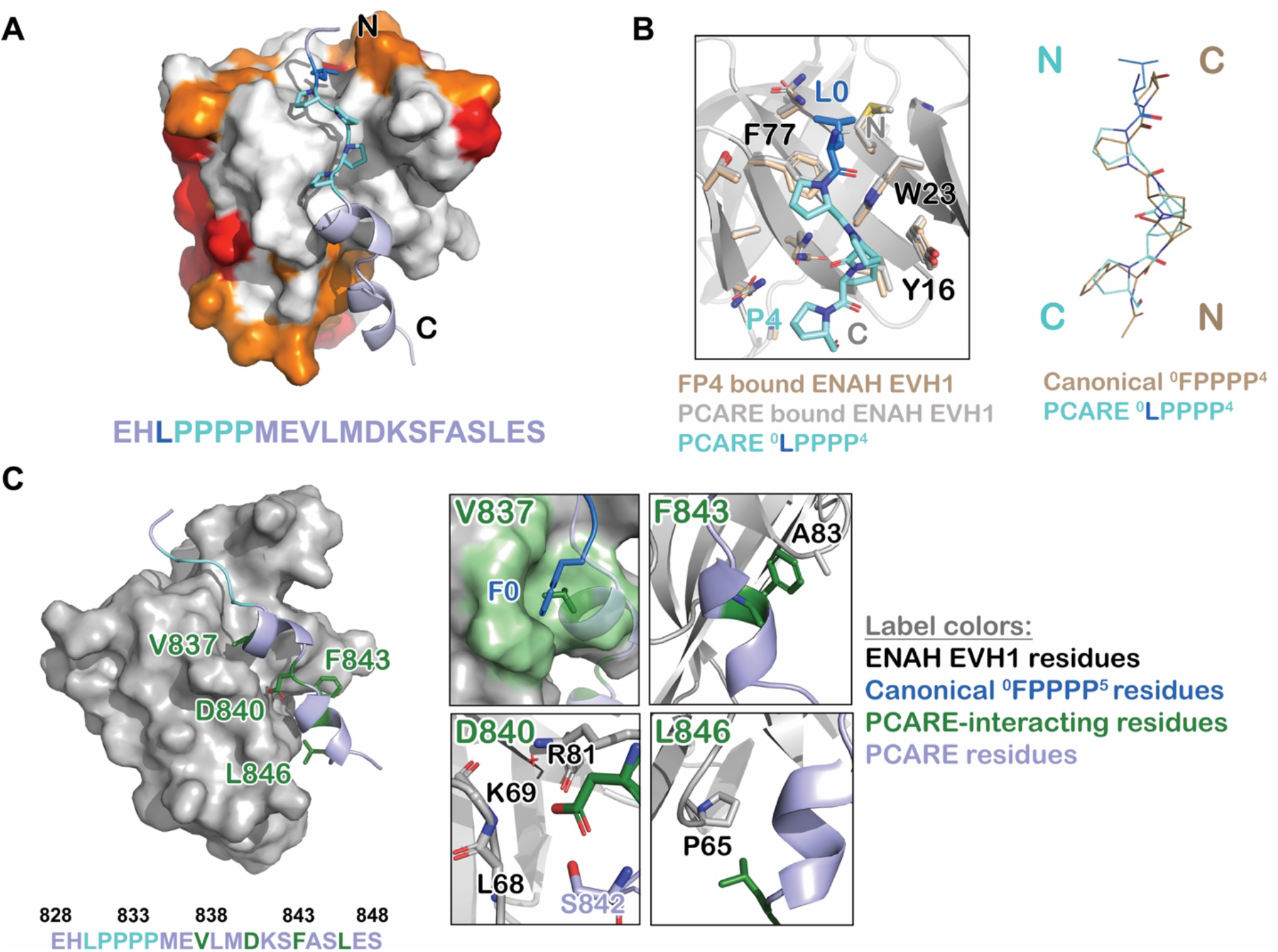
ENAH EVH1 binds to PCARE differently than to the peptide FPPPP. (A) Surface representation of ENAH EVH1 bound to a segment of PCARE, highlighting Ena/VASP EVH1 domain conservation using colors as in Figure 1. (B) View comparing the orientations of an FP_4_ peptide and the LP_4_ region of PCARE. Side chains of the ENAH EVH1 domain are shown as sticks using tan for the FP_4_ complex and grey for the PCARE complex. (C) Surface representation of ENAH EVH1 domain bound to the PCARE peptide. The LP_4_ residues are light blue and other EVH1-interacting residues are green; insets show details of the interactions. Note that the PCARE^828-848^ peptide is numbered as 133-153 in the PDB file.

Contacts between the extended, alpha-helical region of PCARE and ENAH are shown in Figure 3C. PCARE residues Phe843 and Leu846 make hydrophobic interactions with Ala83 and Pro65 on ENAH. The side chain of Asp840 on PCARE docks into a polar pocket on ENAH made up of the backbone atoms of ENAH residues Lys69 and Arg81. Notably, the backbone NH and sidechain hydroxyl group of Ser842 on PCARE form hydrogen bonds with the side chain of Asp840, positioning Asp840 to hydrogen bond with a water molecule that is further coordinated by the backbone NH of ENAH Arg81. Most intriguing is the interaction between Val837 on PCARE and the hydrophobic groove in ENAH, which typically engages large aromatic residues. The alpha-helical structure of PCARE^828-848^ buries the smaller Val837 in the same site where phenylalanine can bind. Collectively, our results reveal a noncanonical mode of binding where the FP_4_ motif of PCARE binds to the ENAH EVH1 domain in a reversed N-to-C orientation and makes extra contacts to achieve high affinity.

### PCARE achieves paralog selectivity by stabilizing an ENAH EVH1 domain-specific conformation with a novel FP_4_ flanking sequence element

Interestingly, our structure of ENAH EVH1 domain bound to PCARE^828-848^ shows that 16 of the 18 residues that are within 4 Å of PCARE in ENAH are identical in VASP and EVL. Thus, whereas the FP_4_-binding site is 100% conserved, the extended binding site engaged by PCARE is also highly conserved. Residue 63 is alanine in ENAH and VASP, and the corresponding residue 64 in EVL is serine. Modeling serine at position 63 in the PCARE-bound structure of ENAH shows that the side-chain hydroxyl group can be readily accommodated in a solvent-facing conformation without interfering with PCARE binding. On the other hand, residue 65 is proline in ENAH and the corresponding residue is valine in EVL and VASP. ENAH Pro65 makes extensive contacts with PCARE and is largely buried at the domain-peptide interface (Figure 3C). We speculated that proline vs. valine might contribute to the ENAH-binding preference of PCARE. To test this, we made EVL with a valine-to-proline mutation, with the expectation that this would increase PCARE B binding affinity. Surprisingly, EVL V65P bound to PCARE B 5-fold weaker than did wild-type EVL (K_D_ = 112.1 μM vs. 22 μM, Table 1). In contrast, EVL V65P bound to an FP_4_-containing ActA peptide, which does not contact Val65 (Barone et al., 2020), with the same affinity as wild-type EVL (K_D_ = 2.4 μM vs. 2.7 μM, Table 1), indicating that the V65P mutation does not lead to global disruption of the domain structure.

**Table 1.**
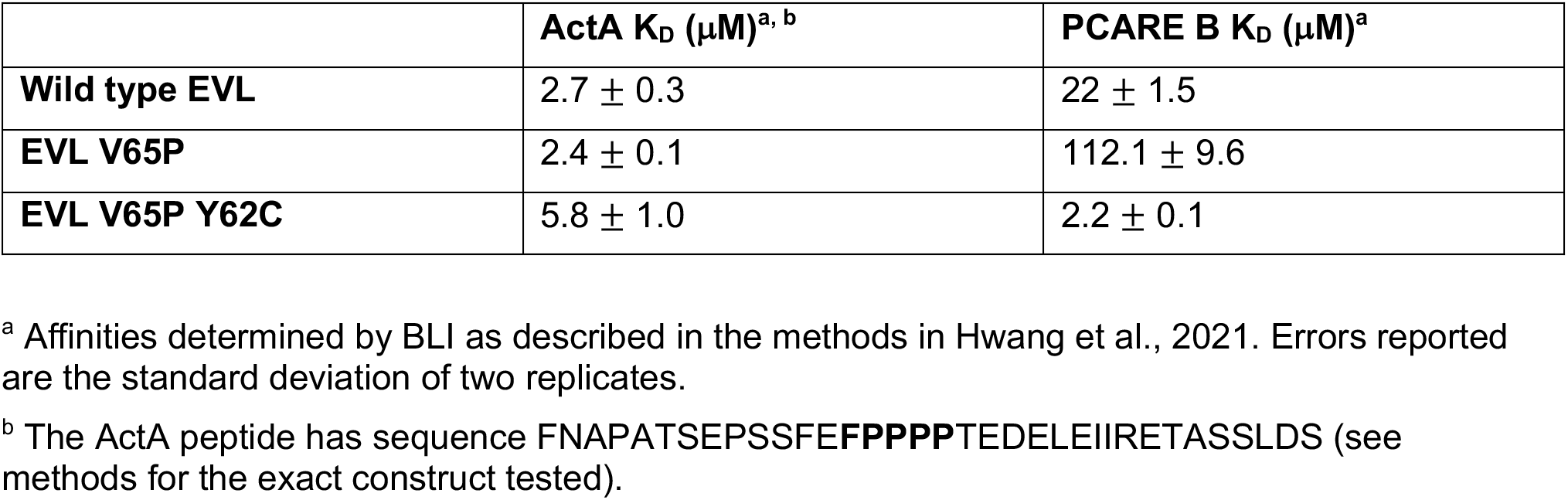
Affinity of EVL EVH1 domain mutants for ActA peptide and PCARE B.

Comparing the structures of ENAH EVH1 domain bound to PCARE^828-848^ vs. the peptide FPPPP (PDB 1EVH; Prehoda et al., 1999) shows a conformational difference in ENAH: a loop composed of residues 80-86, which forms part of the extended PCARE binding site, is shifted by 3 Å (Figure 4A). Structure gazing suggested that hydrophobic core residues Tyr63 in EVL (Cys62 in ENAH), and Trp89 and Leu15 in VASP (Tyr87 and Val15 in ENAH), are incompatible with this conformational change (Figure 4A). To test this, we made EVL V65P Y62C. This EVL double mutant bound to PCARE B with K_D_ = 2.2 μM, which is 56-fold lower than the K_D_ for binding to EVL V65P (Table 1). This striking enhancement in affinity indicates strong coupling between these two mutated positions in EVL. However, these two mutations alone enhanced binding to EVL by only 10-fold over wild type, whereas the difference in binding affinity between ENAH and EVL is 70-fold (Figure 1C). We concluded that a broader set of residues must contribute to stabilizing the ENAH-specific conformation, but it was not readily apparent which residues these might be.

**Figure 4.**
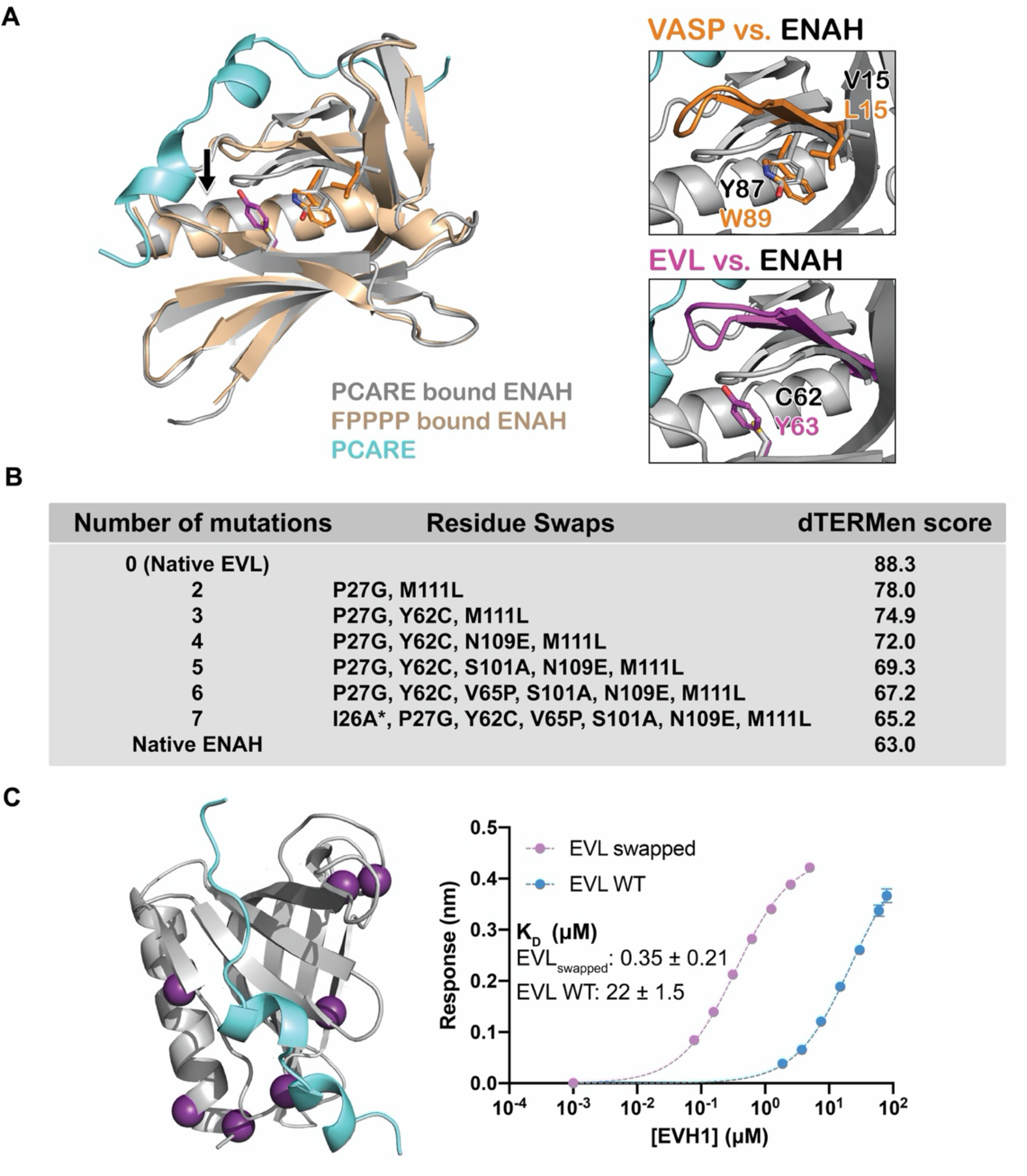
PCARE achieves paralog selectivity by stabilizing an ENAH EVH1 domain-specific conformation. (A) Superposition of ENAH EVH1 domains bound to PCARE or FP_4_ peptide (1EVH). The black arrow highlights a 3 Å shift in a loop that forms part of the binding pocket. Insets show residues that differ between ENAH and VASP or EVL near this loop. (B) Lowest dTERMen energy obtained when swapping 0-6 residues from ENAH into EVL, when modeling on the structure of ENAH EVH1 bound to PCARE. * indicates the mutation was added based on manual inspection. (C) ENAH EVH1 domain bound to PCARE peptide, with residues that were swapped into the EVL EVH1 domain to rescue affinity marked as purple spheres. On the right are binding curves for WT EVL EVH1 domain and EVL_swapped_ EVH1 domain binding to PCARE B. Error reported as the standard deviation of two replicates.

To identify the ENAH residues responsible for PCARE binding specificity, we used the structurebased modeling method dTERMen (Zhou et al., 2017). dTERMen is a protocol for scoring the compatibility of a sequence with a backbone structure. Energies are computed based on the frequencies with which combinations of residues are found in tertiary motifs in known protein structures. As expected, when scoring different sequences on the structure of ENAH bound to PCARE, the EVL sequence scored considerably worse than the sequence of ENAH itself. Guided by the dTERMen score, we introduced increasing numbers of residues from ENAH into EVL. Seven replacements were sufficient to recapitulate dTERMen energies similar to that for ENAH in the PCARE^828-848^-bound conformation (Figure 4B). These residues are distributed across the EVH1 domain, and several are distant from the PCARE^828-848^ binding site (Figure 4C). We made a mutated EVL EVH1 domain with the 7 corresponding residues from ENAH. This protein, EVL_swapped_, bound as tightly to PCARE B as did ENAH EVH1 (K_D_ = 0.35 vs. 0.32 μM) (Figure 4C). Given that wild-type EVL and ENAH differ at 29 sites, and there are 1.56 million potential residue swaps of 7 residues, it is particularly notable that dTERMen quickly led us, in just a single attempt, to a combination of residues sufficient to transfer binding specificity.

### Engineered binders engage ENAH EVH1 domain with increased affinity and specificity

Based on our structure of PCARE bound to ENAH, we aimed to design even higher affinity, ENAH-selective peptides. To this end, we took a rational design approach. Our strategy relied on designing peptides that can simultaneously engage two binding sites on ENAH EVH1: the canonical FP4-binding site that is occupied by FP4 peptides such as ActA, ABI1, and PCARE (Prehoda et al., 1999; Federov et al., 1999, Barone et al., 2020; Hwang et al., 2021), and a noncanonical site previously identified in VASP EVH1 that we have shown is also important for certain ENAH-peptide complexes (Hwang et al., 2021; Acevedo et al., 2017) (Figure 5A, B). Using a structural model based on PDB structure 5NC7, we estimated appropriate lengths for connecting linkers (Hwang et al., 2021; Barone et al., 2020). We then made and tested different combinations of binding motifs and linkers that we predicted could bridge these two sites.

**Figure 5.**
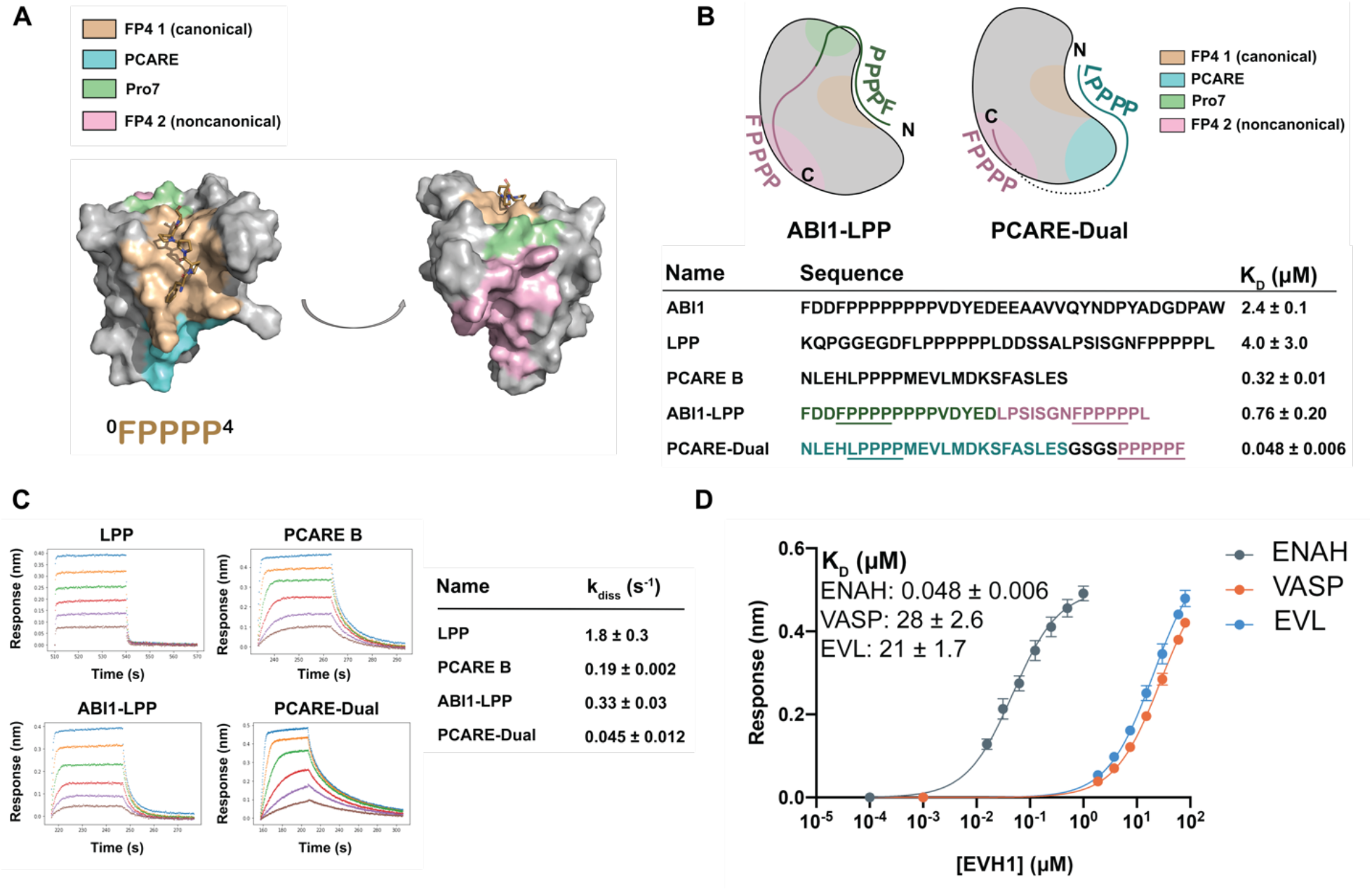
Engineered peptides bind to the ENAH EVH1 domain with high affinity and specificity. (A) Surface representation of ENAH EVH1 with binding sites discussed in this study indicated. (B) Design scheme for high affinity binders ABI1-LPP and PCARE-Dual. (C) BLI binding and dissociation curves. Blue, orange, green, red, purple, and brown curves denote EVH1 concentrations in descending order. LPP: 80, 36, 16, 7.0, 3.1, 1.4 μM. PCARE B and ABI1-LPP: 2.5, 1.3, 0.63, 0.31, 0.16, 0.078 μM. PCARE-Dual: 0.50, 0.25, 0.0625, 0.031, 0.016, 0.0078 μM. Values reported as k_diss_ ±SD for two independent BLI replicates. (D) BLI binding curves for PCARE-Dual binding to the EVH1 domains of ENAH, VASP, or EVL. Errors for 5B and 5D are reported as the standard deviation of two replicates.

In one design, we fused the high-affinity PCARE B sequence, via a short GSGS linker, to a second FP4 motif designed to support bivalent binding while stabilizing the ENAH-specific conformation. Peptide PCARE-Dual bound ~7-fold tighter than PCARE B, with K_D_ = 50 nM (Figure 5B, D). In a second design, we used a peptide from ABI1 in place of PCARE B and fused this to a peptide from the protein LPP. ABI1 is an Ena/VASP interaction partner that contains the sequence FP8. We have shown that proline residues C-terminal to the FP4 motif, as well as surrounding acidic residues, enhance affinity for the ENAH EVH1 domain (Hwang et al., 2021). In particular, the seventh proline of the ABI1 FP4 motif engages ENAH EVH1 at what we call the Pro7 site (green in Figures 5A and B). LPP is another ENAH binding partner that we have shown engages the noncanonical EVH1 binding site. We fused a 17-residue segment of ABI1 to part of the LPP linker and a second FP4 motif to make ABI1-LPP (Figure 5B). Our rationale was that the ABI1-derived segment would occupy the canonical FP4 binding site and the LPP linker would wrap along the surface of the EVH1 domain and position a second FP4 motif near the noncanonical FP4 site. The ABI1-LPP fusion peptide bound with K_D_ = 0.76 μM, which is 3-fold tighter than the ABI1 portion alone and 5-fold tighter than the LPP portion alone, although weaker overall than PCARE-Dual.

The enhancements in affinity for PCARE-Dual and ABI1-LPP come from decreases in off-rate, with PCARE-Dual dissociating ~100-fold slower than dual-motif peptide LPP (Figure 5C). Finally, we found that PCARE-Dual is 400-600-fold selective for ENAH over EVL and VASP, even though VASP is also known to have a noncanonical binding site that can engage a second FP4 motif (Acevedo et al., 2017), providing the tightest and most specific known binder to the ENAH EVH1 domain to date (Figure 5D).

## Discussion

Although the three paralogs have some distinct cellular functions, proteins ENAH, VASP, and EVL are highly conserved in sequence and structure. The EVH1 domains are 100% identical in sequence in the core FP4-binding groove and share 62-72%sequence identity through the rest of the EVH1 domain (Figure 1A, B). FP4-motif peptides engage a small and relatively flat surface on EVH1; previously solved structures have shown how short peptides bind to this core region or, in some cases, to an immediately adjacent hydrophobic patch (Barone et al., 2020). The limited contacts with a highly conserved, shallow site make it challenging to achieve high-affinity or paralog-selective binding.

There are currently two proteins reported to bind specifically to the ENAH EVH1 domain. The LIM domain of testin binds to ENAH EVH1 with a K_D_ of 3 μM and does not bind detectably to the EVL or VASP EVH1 domains. Testin achieves paralog specificity by making contacts with ENAH outside of the canonical FP4 groove, at surface sites where the paralogs differ (Boëda et al., 2007). Synthetic mini-protein pGolemi binds to ENAH with K_D_ = 0.29 μM, moderate selectivity (20-fold) over VASP, and higher selectivity (≥120-fold) over EVL (Golemi-Kotra et al., 2004). The mechanism behind the specificity of pGolemi has not been determined, and mutational data do not readily rationalize a model (Holtzman et al., 2007).

We have now shown that the protein PCARE contains residues adjacent to an FP4 motif that confer both high affinity and selectivity for ENAH over VASP and EVL by stabilizing an ENAH-specific conformation. The biological implications of the specificity of PCARE towards ENAH are unclear at this point. ENAH, VASP, and EVL induce distinct actin remodeling phenotypes based on their differential preferences for profilin isoforms (Mouneimne et al., 2012). Thus, paralog selectivity could be necessary when paralog-dependent actin dynamics and structures are required. PCARE is expressed in retinal photoreceptor cells and interacts with actin modulator WASF3 (Corral-Serrano et al., 2020). This interaction localizes actin remodeling to the base of the outer segment, a highly modified primary cilium, to help initiate new outer segment disk formation. Our results suggest that ENAH, but not VASP or EVL, may promote this process.

A dramatic feature of the PCARE-bound ENAH structure is that the LPPPP motif of PCARE binds in a reversed orientation, relative to previously observed FP4 ligands. The PPII helix possesses two-fold rotational pseudosymmetry, meaning that the side chains and backbone carbonyls are similarly positioned in either the N-to-C- or C-to-N-terminal directions. SH3 domains exploit this pseudosymmetry to bind proline-rich sequences in either direction (Zarrinpar et al., 2003). Although all Ena/VASP and Homer EVH1 domain structures solved so far show the PPII helix engaged in a single direction, the WASP EVH1 domain binds its proline-rich ligand in the opposite direction (Volkman et al., 2002). Consequently, it has long been hypothesized, although never demonstrated, that Ena/VASP EVH1 domains might also bind FP4 motifs in either direction (Ball et al., 2002). Here we show that this is the case for ENAH. Interestingly, the large effect on PCARE binding that we observed for EVL V65P compared to wild-type EVL EVH1 suggests that EVL also binds PCARE in a reversed orientation (relative to FP4 ligands such as ActA), albeit weakly, because this is the orientation that best explains contacts with position 65.

We are not aware of other examples of such dramatic conformational specificity, in which a natural ligand binds selectively to one paralog despite almost complete (~89%) conservation of the binding site amongst its family members. However, this mechanism is reminiscent of how the cancer drug Gleevec achieves 3000-fold selectivity for Abl over Src, despite the fact that the two proteins share ~46% sequence identity across the kinase domain, and ~86% identity in the Gleevec binding site (Seeliger et al., 2007). The mechanism for the selectivity of Gleevec long eluded explanation. As we found here, a few residue swaps based on sequence alignments and structure gazing failed to rescue the affinity of Src for Gleevec, because these mutations failed to account for the higher-order epistatic interactions that contribute to specificity (Seeliger et al., 2007). Interestingly, because evolution has sampled kinase sequence space under the constraints of epistasis, retracing the evolutionary trajectories that led to extant Src and Abl kinases, using ancestral reconstruction, allowed Wilson et al. to identify residues involved in a hydrogen-bonding network distant from the Gleevec binding interface that are key for selective binding to Abl (Wilson et al., 2015).

We used an alternative approach to discover a set of residues that contribute to specificity, turning to structure-based modeling. Using dTERMen, we scored the compatibility of different sequences with the ENAH-PCARE^828-848^ structure template and identified residues in EVL that are incompatible with this binding mode. Our method solved the challenging problem of identifying mutations within an epistatic network that contribute to function. Importantly, this facile method is easily generalizable to many protein systems as long as a structure is available.

Our results demonstrate an intriguing example of how nature has evolved a protein that achieves selectivity by exploiting a conformation accessible to only one paralog, rather than by making contacts with paralog-specific residues. Our findings raise the question of how widely this mechanism of selectivity is exploited by other paralogous families, especially within modular interaction domain families. As evidenced by residues identified by dTERMen in our residue swap experiments, hydrophobic core packing plays a key role in enabling new ligand-bound conformations. A recent study demonstrated that randomizing hydrophobic core residues of the SH3 domain from human Fyn tyrosine kinase could effectively switch its affinity and specificity to different ligands (Ben-David et al., 2018). In addition, directed evolution experiments that varied only the hydrophobic core residues of ubiquitin yielded a conformationally stabilized ubiquitin variant that was specific for the deubiquitinase USP7 (Zhang et al., 2012). These results hint that such mechanisms of conformational specificity could be widespread.

The PCARE peptide identified in this study will serve as a valuable research tool to dissect ENAH-specific biological functions in cells. Cytoplasmic expression of a PCARE peptide effectively displaces ENAH from focal adhesions but not membrane protrusions, indicating that this peptide could serve as a valuable tool to selectively perturb EVH1-dependent functions of ENAH. In contrast, a mito-tagged PCARE peptide selectively sequesters ENAH at the mitochondria, depleting it from sites of normal function and inhibiting ENAH-dependent biological processes such as cell adhesion. This is likely due to the high avidity of these peptides when localized to the mitochondrial surface, which could potentially outcompete interactions between the EVH2 domain of ENAH and actin-based structures (Bachmann et al., 1999).

There is emerging evidence in the literature that paralog-specific functions exist within the Ena/VASP family. For example, ENAH, but not VASP or EVL, is a regulator of local translation in neurons (Vidaki et al., 2017). In contrast, EVL has a specific role in durotactic invasion (Puleo et al., 2019). Our mito-tagged peptides will serve as valuable research reagents in uncovering and characterizing ENAH-specific molecular functions in other tissues, such as the retina.

The role of ENAH in cancer metastasis has motivated work to identify inhibitors of its EVH1-mediated interactions, and extensive structure-based design and chemical optimization recently led to a high-affinity molecule (K_D_ = 120 nM) that reduced breast cancer cell extravasation in a zebrafish model. This compound and its analogs mimic the PPII conformation of an FP4 motif. However, because the molecule binds to the highly conserved FP4-binding site, this inhibitor has similar affinity for ENAH, EVL, and VASP (Barone et al., 2020). Paralog-specific Ena/VASP inhibitors could have higher therapeutic potential, given the antagonistic roles of ENAH and EVL in promoting and suppressing breast cancer metastasis. ENAH deficiency in mouse models of cancer has been shown to decrease metastasis by reducing tumor cell invasion and intravasation (Roussos et al., 2011). On the other hand, EVL is known to promote the formation of cortical actin bundles that suppress the invasion of breast cancer cells (Padilla-Rodriguez et al., 2018). Thus, developing high affinity, paralog-specific inhibitors of Ena/VASP proteins could be crucial in reducing pleiotropic side effects when treating metastasis. The PCARE B peptide exhibits exquisite affinity and specificity for ENAH. We envision it, as well as our engineered PCARE-Dual binder, as a promising lead for developing therapies to treat ENAH-dependent diseases.

## Methods

### Protein expression and purification

Monomeric ENAH, EVL, or VASP EVH1 domains for use in BLI and ITC experiments, and ENAH EVH1-peptide fusions for crystallography were cloned into a pMCSG7 vector (gift from Frank Gertler, MIT), which places a 6x-His and TEV cleavage tag N-terminal to the EVH1 domain. These constructs were transformed into Rosetta2(DE3) cells and grown in 2xYT media supplemented with 100 μg/mL ampicillin. Cells were grown while shaking at 37 °C to an O.D. 600 of 0.5-0.7 and then cooled on ice for at least 20 minutes. Cells were then induced with 0.5 mM IPTG and grown while shaking at 18 °C overnight. Induced cultures were resuspended in 25 mL of wash buffer (20 mM HEPES pH 7.6, 500 mM NaCl, 20 mM imidazole) and frozen at −80 °C overnight. The next day, cultures were sonicated, spun down, and applied to Ni-NTA agarose resin equilibrated with wash buffer, and washed as described above. Samples were eluted in 10 mL elution buffer (20 mM HEPES pH 7.6, 500 mM NaCl, 30 mM imidazole). TEV protease was added to the elution at a ratio of 1 mg TEV:50 mg tagged protein along with 1 mM DTT. This mixture was dialyzed against TEV cleavage buffer (50 mM HEPES pH 8.0, 300 mM NaCl, 5 mM DTT, 1 mM EDTA) at 4 °C overnight and then applied over Ni-NTA agarose resin equilibrated with wash buffer. The column was washed with 2 x 8 mL of wash buffer, and the resulting flow-through and washes were pooled, concentrated, and applied to an S75 26/60 column equilibrated in gel filtration buffer (20 mM HEPES pH 7.6, 150 mM NaCl, 1 mM DTT, 1% glycerol). Purity was verified by SDS-PAGE and combined fractions were concentrated and flash frozen at −80 °C.

### Small-scale protein purification for biolayer interferometry

SUMO-peptide fusions were cloned into a pDW363 vector that appends a BAP sequence and 6x-His tag to the N-terminus of the protein and transformed into Rosetta2(DE3) cells (Novagen). Rosetta2(DE3) cells encoding SUMO-peptide fusions were grown in 20 mL of LB + 100 μg/mL Ampicillin + 0.05 mM D-(+)-Biotin dissolved in DMSO for *in vivo* biotinylation. Cells were grown to an OD of 0.5-0.7 with 1 mM IPTG and induced for 4-6 hours at 37 °C. Pellets were spun down and frozen at −80 °C for at least 2 hours. Pellets were then thawed and resuspended in B-PER reagent (ThermoFisher) at 4 mL/gram of pellet with 0.2 mM PMSF. This suspension was shaken at 25 °C for 10-15 minutes and then spun down at 15,000 g for 10 minutes. The supernatant was applied to 250 μL of Ni-NTA agarose resin equilibrated in 20 mM Tris pH 8.0, 500 mM NaCl, 20 mM Imidazole (Buffer A), and then washed 3 times with 1 mL of this buffer. Peptides were eluted in 1.8 mL of 20 mM Tris pH 8.0, 500 mM NaCl, 300 mM imidazole to use in BLI assays.

### Biolayer interferometry (BLI)

All BLI experiments were performed with 2 replicates on an Octet Red96 instrument (ForteBio). Biotinylated, 6x-His-SUMO-peptide fusions purified in small scale were diluted into BLI buffer (PBS pH 7.4, 1% BSA, 0.1% Tween-20, 1 mM DTT) and immobilized onto streptavidin-coated tips (ForteBio) until loading reached a response level between 0.5-0.6 nm. The loaded tips were immersed in a solution of ENAH EVH1 domain at a relevant dilution series in BLI buffer at an orbital shake speed of 1000 rpm and data were collected until the binding signal plateaued. ENAH-bound tips were subsequently placed into BLI buffer for dissociation and data were collected until the binding signal plateaued. K_D_ values were obtained through steady-state analysis. Briefly, the data were corrected for background by subtracting the signal obtained when doing the same experiment using biotinylated, 6x-His-SUMO lacking any peptide, instead of an immobilized ENAH-binding peptide. The association phases were then fit to a one-phase binding model in Prism and the equilibrium steady-state binding signal values from that fit were plotted against ENAH concentration and fit to a single-site binding model in Prism to obtain dissociation constants. Errors are reported as the standard deviation of two replicates.

### Crystallography

Crystals of ENAH fused at the C-terminus to PCARE were grown in hanging drops containing 0.1M Tris pH 8.0 and 3.30 M NaCl at 18 °C. 1.5 μL of ENAH-PCARE (769 μM in 20 mM HEPES, 150 mM NaCl, 1 mM DTT) was mixed with 0.5 μL of reservoir solution, and football-shaped crystals appeared in two days. Diffraction data were collected at the Advanced Photon Source at Argonne National Laboratory, NE-CAT beamline 24-IDE. The ENAH-PCARE data were integrated and scaled to 1.65 Å with XDS, and the structure was solved with molecular replacement using the ENAH EVH1 structure 6RD2 as a search model. The structure was refined with iterative rounds of model rebuilding with PHENIX and COOT. Table S1 reports refinement statistics. The structure is deposited in the PDB with the identifier 7LXF. Note that the PCARE^828-848^ peptide is numbered as 133-153 in the PDB file in accordance with the ENAH-PCARE fusion protein numbering.

### Modeling using dTERMen

The dTERMen scoring function and protocol are described in Zhou et al., 2020. The method requires that a template-specific scoring function be computed, based on statistics derived from structures in the PDB. After that, the inputs to dTERMen are the backbone coordinates of a structure and a sequence. dTERMen returns a score for the input sequence adopting the input structure; lower scores correspond to lower energies. Side-chain positions are not modeled explicitly. To score the EVL sequence on the ENAH-PCARE backbone template, we generated pairwise alignments of the EVH1 domains of ENAH and EVL to determine how to map sequence to structure. The EVL EVH1 domain is longer than that of ENAH by one residue, so Lys27 was removed from EVL. Note that Lys27 was also removed from the manually chosen EVL V65P and EVL V65P Y62C mutants.

We then used dTERMen to score all possible combinations of residue swaps between EVL and ENAH, up to 6 possible positions. Residue swap combinations that led to the minimum energy score were recorded. The best 6 mutations were sufficient to nearly recapitulate the energy score of the native ENAH sequence on the ENAH PCARE template. We also included an I26A mutation based on manual inspection of the ENAH-PCARE^828-848^ structure, which also lowered the dTERMen energy. We cloned, overexpressed, and purified this swapped EVL sequence, as described in Hwang et al., 2021, to test for binding to PCARE B

### Plasmids for cell culture

For experiments in mammalian cells, the following plasmids were used: pLKO.1-Enah shRNA (GE Dharmacon TRCN0000061827, *Homo sapiens* antisense 5’- TTAGAGGAGTCTCAACAGAGG -3’), pLKO.1-Evl shRNA (Sigma TRCN0000091075, *Mus musculus* antisense 5’-TTGTTCATTTCTTCCATGAGG-3’), and non-targeting pLKO.1 control (a gift from Felicia Goodrum, University of Arizona). GFP, and mouse cDNA sequences for GFP-tagged ENAH, VASP, and EVL (gifts from Frank Gertler, MIT) were sub-cloned into the pCIB lentiviral expression vector (Addgene plasmid #120862) as previously described (Puleo et al. 2019). ENAH EVH1 domain deletion mutant was generated using inverse PCR site-directed mutagenesis of full-length ENAH. All sequences of constructed plasmids were confirmed by Sanger sequencing. mRuby2-Pcare and Mito-mRuby2-Pcare inserts were synthesized (Twist Bioscience) and subcloned into a SFFV-promoter lentiviral expression vector.

### Cell culture

MCF7 and HEK293T cells were cultured in high-glucose Dulbecco’s modified Eagle’s medium (DMEM) base media with sodium pyruvate (Corning) supplemented with 10% fetal bovine serum (FBS; Millipore), 2 mM L-glutamine (Corning), and 100 U/mL penicillin with 100 μg/mL streptomycin (Corning). MV^D7^ cells were cultured in high-glucose DMEM supplemented with 15% FBS, 2 mM L-glutamine, 100 U/mL mouse interferon-γ (Millipore). MCF7 and HEK293T cells were maintained in a 37 °C humidified incubator under 5% CO_2_. MV^D7^ cells were maintained in a 32 °C humidified incubator under 5% CO_2_. For lentiviral production, second-generation lentiviral particles were generated by PEI transfection of 293T cells as previously described (Yang et al., 2017) with transfer plasmid, pMD2.G, and psPAX2 (Addgene #12259, #12260, gifts from Didier Trono). HEK293T media containing lentiviral particles was collected, filtered, and added directly to cultures with polybrene (Gibco). Puromycin (2 μg/mL final concentration; Thermo Fisher) and blasticidin (4 μg/mL final; Gibco) were used to select for cells stably expressing shRNA sequences or Ena/VASP constructs, respectively, after lentiviral transduction. For immunofluorescence experiments, MCF7 cells were cultured on fibronectin-coated coverslips (10 μg/mL; Corning). For live-cell imaging (MV^D7^), cells were cultured in fibronectin-coated glass-bottom dishes (Mattek).

### Reverse-transcription quantitative PCR (RT-qPCR)

To assess *EVL* knockdown in MV^D7^, total cellular RNA was isolated using the Isolate II RNA kit (Bioline) according to the manufacturer’s instructions. cDNA was synthesized from 1000 ng of input RNA using qScript cDNA Synthesis kit (Quantabio). RT-qPCR reactions were run in duplicate on an ABI 7500 Fast Real-Time PCR System (Applied Biosystems) with PowerTrack SYBR Green Master Mix (Thermo Fisher). Primer pairs were confirmed to have 85–110% efficiency based on the slope of the standard curve from a cDNA dilution series. C_T_S were normalized to the C_T_ *GAPDH* housekeeping genes. Percent knockdown was determined using the comparative C_T_ method. *Mus musculus GAPDH* Fwd AGGTCGGTGTGAACGGATTTG, Rev GGGGTCGTTGATGGCAACA. *EVL* Fwd TGAGAGCCAAACGGAAGACC, Rev TTCTGGACAGCAACGAGGAC.

### Western blotting

Cells were lysed in buffer containing 140 mM NaCl, 10 mM Tris pH 8.0, 1 mM EDTA, 0.5 mM EGTA, 1% Triton X-100, 0.1% sodium deoxycholate, and 0.1% SDS with protease and phosphatase inhibitors (Boston Bio Products). Samples were resolved by SDS-PAGE and transferred onto nitrocellulose membranes. Membranes were blocked in Odyssey Blocking Buffer (LI-COR) for 1 hour and incubated at 4 °C overnight with primary antibodies. Primary antibodies were used as follows: mouse Actin 1:2500 (ProteinTech Group, 66009-1-Ig), mouse GAPDH 1:1000 (Cell Signaling Technology, 5174S), rabbit ENAH 1:250 (Sigma, HPA028696), rabbit EVL 1:1000 (Sigma, HPA018849), rabbit VASP 1:1000 (Cell Signaling Technology, 3132S). Membranes were incubated with secondary antibodies conjugated to either Alexa Fluor 680 or 790 (Thermo Fisher) for 1 hour. Immunoblots were scanned using Odyssey CLx imager (LI-COR).

### Immunofluorescence

MCF7 cells were fixed and immunolabeled 8 hours after plating onto fibronectin (10 μg/mL; Corning) coated coverslips to assay focal adhesions. Cells were fixed with 4% paraformaldehyde (PFA; Electron Microscopy Services) with 0.075 mg/mL saponin (*Alfa Aesar*,Sigma) diluted in phosphate-buffered saline (PBS) at 37 °C for 10 minutes. PFA was quenched with 100 mM glycine in PBS at room temperature for 10 minutes. Cells were then blocked in 1% BSA plus 1% FBS in PBS either overnight at 4 °C or for 1 hour at room temperature. The following immunofluorescence reagents and antibodies were used: mouse anti-Paxillin (1:200, clone: 349; BD Biosciences, 612405), rabbit anti-ENAH (1:100; HPA028448, Sigma), goat antimouse Alexa Fluor 488 and Alexa Fluor 647 (1:1000, Thermo Fisher), donkey anti-rabbit Alexa Fluor 405 (Abcam), and Alexa Fluor 488- and Alexa Fluor 647-Phalloidin (1:40, Thermo Fisher). Primary antibodies were diluted in block solution and incubated for 1.5 hours at room temperature. After washing, coverslips were incubated for 1 hour in secondary antibody solution with fluorescently-labelled phalloidin. Coverslips were mounted using ProLong Gold Antifade (Invitrogen) and allowed to cure for at least 24 hours before imaging.

### Microscopy and image analysis

Cells were imaged on a Ti-E inverted microscope (Nikon), with a 100x Apo TIRF 1.49 NA objective (Nikon) and an ORCA-Flash 4.0 V2 CMOS camera (Hamamatsu). For focal adhesion assessment, cells were imaged with total internal reflection fluorescence (TIRF) microscopy. To increase the depth of imaging for examination of mitochondrial localization, standard widefield fluorescence microscopy was used. Focal adhesion quantification was performed as previously described (Puleo et al., 2019). Briefly, a binary mask was generated for paxillin signal and actin signal, denoting focal adhesions and cell area, respectively. To facilitate semi-automated segmentation of focal adhesions, we generated a sharp, high contrast image of the paxillin and actin channels by the following processing steps: deconvolution using 5 iterations of the Richardson-Lucy algorithm, shading correction using rolling ball, and unsharp masked (NIS Elements). Focal adhesion area was quantified by measuring the paxillin area of each cell within the whole cell area, or by examining individual focal adhesions. Each experimental condition was performed in triplicate and plotted together. Images presented in figures have been lightly processed in NIS Elements, including by applying 2 iterations of the Richardson-Lucy deconvolution algorithm and rolling ball shading correction to reduce background in live cell images.

## Acknowledgments

This project was supported by NIGMS award R01 GM129007 to A.E.K. and by NCI award R01 CA196885-01 to G.M. Part of this work is based upon research conducted at the Northeastern Collaborative Access Team beamlines, which are funded by the National Institute of General Medical Sciences from the National Institutes of Health (P30 GM124165). The Eiger 16M detector on the 24-ID-E beamline is funded by an NIH-ORIP HEI grant (S10OD021527). This research used resources of the Advanced Photon Source, a U.S. Department of Energy (DOE) Office of Science User Facility operated for the DOE Office of Science by Argonne National Laboratory under contract DE-AC02-06CH11357. T.H. was partially supported by NIGMS T32 GM007287 and a fellowship from the Koch Institute for Integrative Cancer Research.

We thank the MIT Structural Biology Core for assistance with X-ray crystallography and the MIT Biophysical Instrumentation Facility for instrumentation resources. We thank L. Backman for help with X-ray data collection. We thank F. Gertler and J. Tadros for constructs. We thank members of the Keating lab, Mouneimne lab, and F. Gertler for their thoughtful input on the manuscript.

## Author Contributions

T.H., S.S.P., G.M., and A.E.K. designed the study. T.H., S.S.P., S.M.H., and M.W.I. performed the experiments. R.A.G. helped with X-ray crystallography experiments. T.H., R.A.G., S.S.P, S.M.H., and M.W.I. analyzed data. T.H. and S.S.P wrote the original draft. A.E.K. and G.M. reviewed and edited the paper. A.E.K. and G.M. provided supervision and funding acquisition. G.M. led and supervised all cell-based experiments, while A.E.K. oversaw all other experimental and computational work reported here.

