## Supplementary Materials for "A distributed residue network permits conformational binding specificity in a conserved family of actin remodelers"

### Table of contents:


1. Protein Sequences
2. Figure S1: **Knockdown of *EVL* in MV$^{D7}$ cells**
3. Table S1: **Refinement table for ENAH-PCARE structure**

**Protein sequences**

Constructs below were cloned into a pMCSG7 backbone (gift from F. Gertler) which encodes an N-terminal 6xHis-TEV site. Amino acid sequences of proteins are as given:

**ENAH EVH1**
MHHHHHHSSGVDLGTENLYFQSNAMSEQSICQARAAVMVYDDANKKWVPAGGSTGF
SRVHIYHHTGNNTFRVVGRKIQDHQVVINCAIPKGLKYNQATQTFHQWRDARQVYGLN
FGSKEDANVFASAMMHALEVL*
**VASP EVH1**
MHHHHHHSSGVDLGTENLYFQSNAMSETVICSSRATVMLYDDGNKRWLPAGTGPQA
FSRVQIYHNPTANSFRVVGRKMQPDQQVVINCAIVRGVKYNQATPNFHQWRDARQV
WGLNFGSKEDAAQFAAGMASALEALE*
**EVL EVH1**
MHHHHHHSSGVDLGTENLYFQSNAMSEQSICQARASVMVYDDTSKKWVPI**K**PGQQG
FSRINIYHNTASSTFRVVGVKLQDQQVVINYSIVKGLKYNQATPTFHQWRDARQVYGL
NFASKEEATTFSNAMLFALNIMNSQE*
**EVL EVH1 V65P**[#]
MHHHHHHSSGVDLGTENLYFQSNAMSEQSICQARASVMVYDDTSKKWVPIPGQQGF
SRINIYHNTASNTFRVVGVKLQDQQVVINYSIPKGLKYNQATPTFHQWRDARQVYGLN
FASKEEATTFSNAMLFALNIM*
**EVL EVH1 V65P Y62C**[#]
MHHHHHHSSGVDLGTENLYFQSNAMSEQSICQARASVMVYDDTSKKWVPIPGQQGF
SRINIYHNTASNTFRVVGVKLQDQQVVINCSIPKGLKYNQATPTFHQWRDARQVYGLN
FASKEEATTFSNAMLFALNIM*
**EVL EVH1 swapped (7-residue swapped)**[#]
MHHHHHHSSGVDLGTENLYFQSNAMSEQSICQARASVMVYDDTSKKWVPAGGQQG
FSRINIYHNTASNTFRVVGVKLQDQQVVINCSIPKGLKYNQATPTFHQWRDARQVYGL
NFASKEEATTFANAMLFALEIL*
**ENAH EVH1-PCARE (for crystallography)**
MHHHHHHSSGVDLGTENLYFQSNAMSEQSICQARAAVMVYDDANKKWVPAGGSTGF
SRVHIYHHTGNNTFRVVGRKIQDHQVVINCAIPKGLKYNQATQTFHQWRDARQVYGLN
FGSKEDANVFASAMMHALEVLGGSGSGAAKSEELSCEMEGNLEHLPPPPMEVLMDK
SFASLES*
**SUMO-ActA**
MAGGLNDIFEAQKIEWHEDTGGSSHHHHHHGSGSGSDSEVNQEAKPEVKPEVKPET
HINLKVSDGSSEIFFKIKKTTPLRRLMEAFAKRQGKEMDSLTFLYDGIEIQADQTPEDLD
MEDNDIIEAHREQIGGGFNAPATSEPSSFEFPPPPTEDELEIIRETASSLDS*
**SUMO-PCARE B**
MAGGLNDIFEAQKIEWHEDTGGSSHHHHHHGSGSGSDSEVNQEAKPEVKPEVKPET
HINLKVSDGSSEIFFKIKKTTPLRRLMEAFAKRQGKEMDSLTFLYDGIEIQADQTPEDLD
MEDNDIIEAHREQIGGSGSGNLEH**LPPPP**MEVLMDKSFASLES

[#] These EVL constructs had a single-residue deletion of WT EVL residue Lys 27, which was removed to make the lengths of ENAH EVH1 and the mutated EVL EVH1 domains equal. In PDB structure 1QC6, Lys27 is at the end of a loop that is disordered and has a high B-factor.

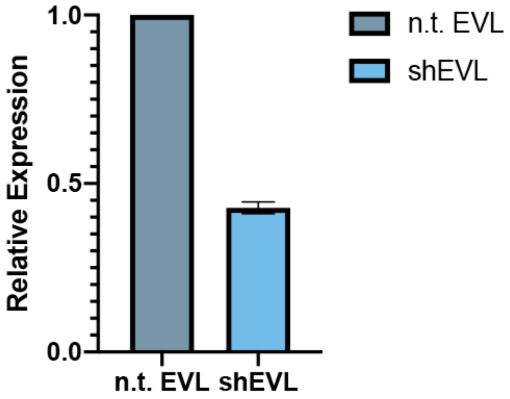


**Figure S1. Knockdown of *EVL* in MV$^{D7}$ cells.** Relative expression of MV$^{D7}_{shEVL}$ versus MV$^{D7}_{ntEVL}$ (nontargeting control) is 42.8 +/- 1.8% by qPCR, 3 biological replicates.

**Table S1. Refinement table for ENAH-PCARE crystal structure**

| | **ENAH-PCARE (7LXF)** |
|---|---|
| **Wavelength** | |
| **Resolution range** | 45.14 - 1.65 (1.71 - 1.65) |
| **Space group** | P 61 2 2 |
| **Unit cell** | 52.12 52.12 197.85 90 90 120 |
| **Total reflections** | 742185 (53799) |
| **Unique reflections** | 20193 (1951) |
| **Multiplicity** | 36.8 (27.6) |
| **Completeness (%)** | 99.87 (99.44) |
| **Mean I/sigma(I)** | 27.78 (0.68) |
| **Wilson B-factor** | 36.47 |
| **R-merge** | 0.0684 (3.906) |
| **R-meas** | 0.0694 (3.979) |
| **R-pim** | 0.0114 (0.750) |
| **CC1/2** | 1 (0.659) |
| **CC*** | 1 (0.891) |
| **Reflections used in refinement** | 20166 (1937) |
| **Reflections used for R-free** | 1012 (98) |
| **R-work** | 0.217 (0.346) |
| **R-free** | 0.235 (0.356) |
| **CC(work)** | 0.960 (0.786) |
| **CC(free)** | 0.936 (0.720) |
| **Number of non-hydrogen atoms** | 1090 |
| **macromolecules** | 1054 |
| **solvent** | 36 |
| **Protein residues** | 134 |

| | |
|---|---|
| **RMS(bonds)** | 0.011 |
| **RMS(angles)** | 1.08 |
| **Ramachandran favored (%)** | 96.15 |
| **Ramachandran allowed (%)** | 3.85 |
| **Ramachandran outliers (%)** | 0.00 |
| **Rotamer outliers (%)** | 0.89 |
| **Clashscore** | 0.48 |
| **Average B-factor** | 48.68 |
| **macromolecules** | 48.85 |
| **solvent** | 43.72 |

Statistics for the highest-resolution shell are shown in parentheses.

Note that the PCARE$^{828\text{-}848}$ peptide is numbered as 133-153 in the PDB file, based on residue numbers in the domain-peptide fusion.
